# Metabolite accumulation from oral NMN supplementation drives aging-specific kidney inflammation

**DOI:** 10.1101/2024.04.09.588624

**Authors:** Tara A. Saleh, Jeremy Whitson, Phoebe Keiser, Praveena Prasad, Brenita C. Jenkins, Tori Sodeinde, Carolyn N. Mann, Peter S. Rabinovitch, Melanie R. McReynolds, Mariya T. Sweetwyne

## Abstract

The mitochondrial-rich renal tubule cells are key regulators of blood homeostasis via excretion and reabsorption of metabolic waste. With age, tubules are subject to increasing mitochondrial dysfunction and declining nicotinamide adenine dinucleotide (NAD^+^) levels, both hampering ATP production efficiency. We tested two mitochondrial interventions in young (6-mo) and aged (26-mo) adult male mice: (ELAM), a tetrapeptide in clinical trials that improves mitochondrial structure and function, and nicotinamide mononucleotide (NMN), an NAD^+^ intermediate and commercially available oral supplement. Kidneys were analyzed from young and aged mice after eight weeks of treatment with ELAM (3 mg/kg/day), NMN (300 mg/kg/day), or from aged mice treated with the two interventions combined (ELAM+NMN). We hypothesized that combining pharmacologic treatments to ameliorate mitochondrial dysfunction and boost NAD^+^ levels, would more effectively reduce kidney aging than either intervention alone. Unexpectedly, in aged kidneys, NMN increased expression of genetic markers of inflammation (IL-1β and Ccl2) and tubule injury (Kim-1). Metabolomics of endpoint sera showed that NMN-treated aged mice had higher circulating levels of uremic toxins than either aged controls or young NMN-treated mice. ELAM+NMN- treated aged mice accumulated uremic toxins like NMN-only aged mice, but reduced IL-1β and Ccl2 kidney mRNA. This suggests that pre-existing mitochondrial dysfunction in aged kidney underlies susceptibility to inflammatory signaling with NMN supplementation in aged, but not young, mice. These findings demonstrate age and tissue dependent effects on downstream metabolic accumulation from NMN and highlight the need for targeted analysis of aged kidneys to assess the safety of anti-aging supplements in older populations.

**Summary Statement:** Declining levels of NAD^+^ and increasing mitochondrial dysfunction with age are functionally linked and are popular mechanistic targets of commercially available anti-aging therapeutics. Studies have focused on nicotinamide mononucleotide (NMN), nicotinamide riboside (NR) and nicotinamide (NAM) supplementation to boost cellular NAD^+^, but a consensus on the dosage and regimen that is beneficial or tolerable has not been reached. We show that although high levels of sustained NMN supplementation are beneficial to liver and heart in aged mice, the same dosing regimen carries age-associated signs of kidney inflammation. Our findings underscore a complex state of age- and tissue-specific metabolic homeostasis and raise questions not only about how much, and for how long, but at what age is NAD^+^ boosting safe.

## Introduction

Advanced age is the single greatest risk factor for kidney disease and injury, with more than 35% of US adults over the age of 65 estimated to have some form of chronic kidney disease (1). Even in older adults without clinically diagnosed kidney disease, decline in kidney function is typical and predictable with age and impacts all cellular segments of the nephron, the ‘functional unit’ of the kidney. The cellular mechanisms underlying this susceptibility are complex, but one mechanism of recent focus is age-associated mitochondrial dysfunction owing to the high mitochondrial endowment of kidneys (2). Mitochondrial dysfunction may be particularly relevant within renal tubule cells, which are highly dependent on mitochondrial activity to generate ATP through fatty acid oxidation and oxidative phosphorylation to support active cellular transport in the processes of blood homeostasis and urine concentration (3–5). This dependence on mitochondrial activity has been successfully exploited as a potential therapeutic target to reduce kidney dysfunction in response to age, disease, and injury (6–8). Furthermore, the levels of NAD^+^, an essential coenzyme that powers these redox reactions, declines with age, and this decline is linked to mitochondrial dysfunction (5). The mechanisms of mitochondrial dysfunction vary and include declining mitochondrial energetic precursors (9), compromised structural integrity (10), and altered turnover/biogenesis (11). Similarly, the modes of action of mitochondrial therapeutic interventions also vary, with two of the most common being those that bolster ATP generation by enhancing the molecular precursor supply and those that improve the integrity or maintenance of the mitochondrial structure.

Two examples of mitochondrial interventions that have been used in both past and current clinical trials are NMN and ELAM. NMN targets NAD^+^ metabolism and contributes to the energetic capacity and cellular sirtuin deacetylase activity (12). Although NMN exists in the body as a natural precursor for NAD^+^, both NMN levels and conversion rates to NAD^+^/NADH fall due to aging. In fact, NMN supplementation restores mitochondrial function (13, 14) by increasing NAD^+^ levels and improving efficiency of the electron transport chain (ETC). ELAM, on the other hand, is a water-soluble, synthetic, four amino acid peptide (D-Arg-2′,6′-dimethyltyrosine-Lys-Phe-NH2), which crosses the cell and mitochondrial membranes and accumulates near the surface of the inner mitochondrial membrane based on electrostatic attraction. ELAM improves the structural integrity of the mitochondrial inner membrane both by altering lipid bilayer packing, and by direct interactions with proteins at the inner membrane (15–17). This structural improvement is thought to cause the improved efficiency of the mitochondrial ETC under ELAM intervention (18–20).

The synergistic effects of these interventions for heart function were first illustrated by our study in 26-month-old mice treated for 8-weeks with combined ELAM and NMN treatment (21). In that study, eight weeks of only ELAM improved diastolic ventricular function, with greater relaxation of the aged ventricle. On the other hand, when treated for eight weeks with only NMN, the aged hearts were better able to respond to dobutamine by pumping harder, demonstrating improvement of systolic function. However, when the treatments were combined, aged mice had simultaneously rejuvenated diastolic and systolic ventricular function, showing that the efficacy of mitochondrial interventions is specific to the underlying cause of dysregulation, which differs based on specific tissue or cell types.

Although we previously demonstrated that late-age mitochondrial intervention with ELAM could reduce age-associated glomerulosclerosis and cellular senescence in the kidneys of C57BL/6J mice (22), similar studies of the kidney with NMN treatment were lacking in aged models. Thus, we aimed to determine whether similar improvements would be found in the kidney in response to either NMN or ELAM+NMN combined treatment and whether these treatments provided additional benefits over ELAM treatment alone. Using the same cohort of mice from the aforementioned heart study, we analyzed the kidneys and livers for markers of cellular senescence, kidney injury and targeted metabolomics. We hypothesized that the NMN and the combined NMN+ELAM treatments reduce or prevent kidney dysfunction in aged mice. Contrary to our hypothesis, and to their effect in young mice, we found that 8 weeks of 300 mg/kg oral NMN supplementation exacerbated renal inflammation pathways in older mice, which were partially reduced by ELAM intervention.

## Results

### Senescence-associated gene expression panel shows Interleukin-1-beta (IL-1β) is upregulated by NMN in the aged kidney, but not liver

ELAM intervention reduces markers of cellular senescence in the aged kidney, and other injury models (22, 23). To determine whether combining NMN with ELAM was similarly, or more, effective, we analyzed gene expression in the kidneys and livers of 26-month-old aged mice with either ELAM, NMN or ELAM+NMN, in addition to young (6-month-old) and aged untreated controls. As senescence cannot be identified through a single gene, a panel of genes known to be involved in senescence signaling were selected for our initial qRT-PCR screen. These included: cell cycle regulators p16 (Cdkn2a), p21 (Cdkn1a) and p53 (Tp53), Lamin B1 (Lmnb1), a nuclear lamina gene known to decrease in response to cellular senescence, and the inflammatory cytokine, IL-1β (24). Other interleukins, including IL8, IL6 and IL-1α were tested but excluded from the panel results as they did not produce quantifiable data by the standard curve.

mRNA expression of IL-1β in kidneys was significantly upregulated in both NMN- (p<0.0001) and ELAM+NMN- (p=0.023) treated aged mice compared to untreated aged mice (Fig 1a). However, in the combined treatment mice, the kidney levels of IL-1β were significantly lower than in NMN treated mice (p=0.0096) (Fig 1a). Expression of IL-1β in kidneys of mice treated with ELAM alone were not different than young or aged untreated animals, suggesting that the upregulation of IL-1β seen with NMN supplementation was ameliorated by the presence of ELAM in the combined treatment (Fig 1a). In liver, the mRNA expression pattern of IL-1β trended opposite that of kidney, with livers from NMN-treated mice having the lowest expression, although no significant changes were detected between treatment or age groups (Fig 1b).

**Figure 1.**
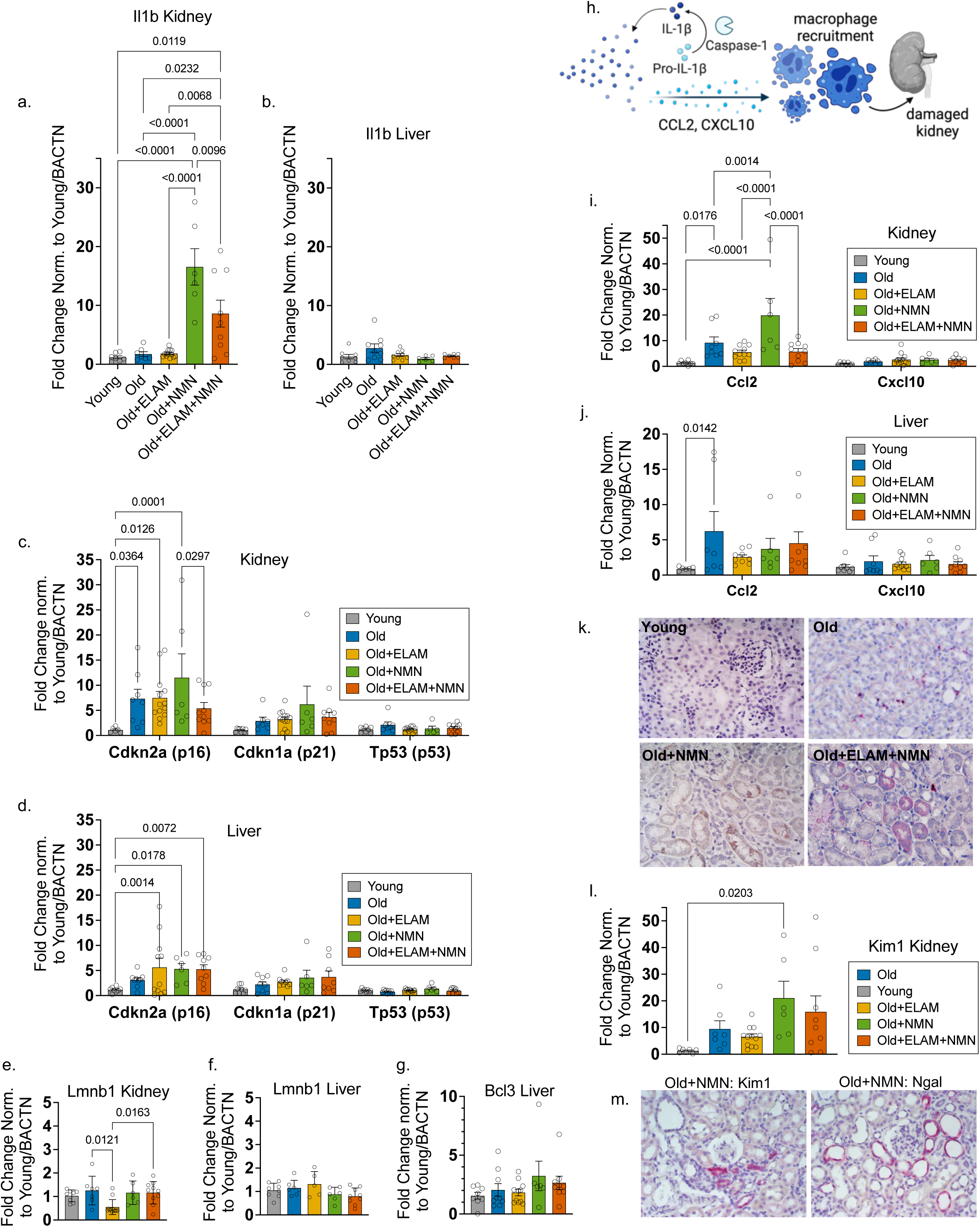
IL-1β and downstream target, CCL2, are differentially regulated by NMN in aged kidneys and livers. mRNA expression levels by quantitative real-time polymerase chain reaction (qRT-PCR) in all figures are from untreated 6-month-old mice or from 26-month-old mice either untreated, or treated with: Elamipretide (ELAM), Nicotinamide mononucleotide (NMN) or ELAM+NMN. Target gene expression is first normalized to housekeeping gene, Bactn, and fold changes are shown relative to the average of young untreated mice. Error = +/-SEM; p-values by 2-way ANOVA with Tukey multivariable testing; sample size (n) indicated with circle symbol for each treatment group. **a&b.** mRNA expression of senescence-associated inflammatory cytokine Interleukin-1-beta (Il1b) in **(a)** kidney. NMN-treated aged animals had statistically significant increase in Il1b kidney expression levels compared to all other groups. Il1b is significantly lower in ELAM+NMN treated kidney compared to NMN-treated kidneys. **(b)** qRT-PCR of Il1b in the liver. **c&d.** mRNA expression of gene targets, Cdkn2a (p16), Cdkn1a (p21), Tp53 (p53) which are known senescence-associated cell cycle regulators in **(c)** the kidney. (**d)** the liver. **e&f.** mRNA expression of senescence-associated biomarker, Lmnb1 (Laminin beta-1), in **(e)** the kidney and **(f)** in the liver. **g.** mRNA expression of Bcl3 (B-cell Lymphoma C) in the liver. **h.** Schematic outlining the mechanism in which IL-1β downstream targets, Ccl2 and Cxcl10, may trigger pro-inflammatory pathway and macrophage recruitment leading to further kidney injury. **i&j.** qRT-PCR panel of mRNA expression for IL-1β downstream targets, Ccl2 and Cxcl10, in (**i**) the kidney and (**j**) the liver. Representative images for RNA In-Situ Hybridization (RNA-ISH). Brown or red punctate dots signal areas of expression of associated gene. **k.** Representative images of RNA In-Situ Hybridization (RNA-ISH) of IL-1β. **l.** qRT-PCR panel of mRNA levels of Kim1 (Kidney Injury Marker-1) in the kidneys. **m.** Representative images of RNA In-Situ Hybridization (RNA-ISH) of Kim1, and Lcn2 (Lipocalin-2/Ngal), a secretory protein associated with distal tubule inflammatory injury.

Similar to the pattern seen in kidney expression levels of IL-1β (Fig 1a), kidney expression of Cdkn2a in ELAM+NMN treated aged mice was significantly reduced by 50% compared to NMN-treated aged mice (mean relative to young fold change: ELAM+NMN 5.40 +/- 2.06 SEM vs. NMN 11.51 +/- 2.15 SEM, p=0.03) (Fig 1c). mRNA expression of Cdkn2a, increased 7-fold (p=0.036) in aged untreated kidneys relative to young untreated but this was not ameliorated by treatment in aged mice with either ELAM or NMN (Fig 1c). Rather, NMN-treated mice showed a trending increase in kidney expression of Cdkn2a relative to aged control, although not reaching significance (Fig 1c). Cdkn2a expression in the kidneys of aged ELAM+NMN-treated group was not significantly different from the young controls but was significantly lower than levels detected in NMN-treated group. Previously, we reported that eight weeks of ELAM treatment reduced senescence in 26-month-old kidneys, as measured by senescence-associated-β-galactosidase, but this reduction was not detected by Cdkn2a expression (22) (Fig 1c). Functioning as a control for kidney-specific versus systemic findings, we also analyzed expression in the liver. Cdkn2a liver expression increased in aged untreated mice but did not reach significance relative to young animals (Fig 1d).

Although Cdkn1a gene expression trended higher in aged tissues than in young, this did not reach significance in either kidney or liver (Fig 1c, d). Changes in expression level of Tp53 between young and aged animals were not detected in either tissue (Fig 1c, d). In the kidney, Lmnb1 levels were significantly decreased in ELAM treated aged mice compared to untreated aged mice (p=0.012) and ELAM+NMN treated mice (p=0.016), in the case of this gene suggesting increased involvement of cellular senescence with exclusive ELAM treatment (Fig 1e). However, Lmnb1 expression was unchanged across treatment and age groups in the liver (Fig 1f). Furthermore, B-cell Lymphoma 3 (Bcl3), an additional hepatic senescence marker, was examined to assess any further liver-specific senescence-related response from these treatments (25). However, there was no significant increase in Bcl3 expression with age or treatment (Fig 1g). While a single marker of senescence cannot be definitive, coupled with earlier results (Fig 1b, d, f), this suggests that the treatments do not impact cellular senescence in the liver.

Considering this organ- and age- specific treatment response, it is important to note the role of IL-1β as a fundamental marker of age-associated inflammation and a component of the senescence-associated secretory phenotype (SASP) (26). In the kidney, increased IL-1β signaling is detected in patients who have experienced an acute kidney injury or who have chronic kidney disease (27, 28). We asked if the increased expression of IL-1β in the kidneys of NMN-treated mice was an indication of increased IL-1β signaling by assaying expression of two of its downstream targets Ccl2, an inflammatory cytokine, and Cxcl10, an inflammatory chemokine (29–32) (Fig 1h).

In kidney, the mRNA expression pattern of Ccl2 was like that of IL-1β with expression significantly higher in NMN-treated mice as compared to any other group (Fig 1i). As with IL-1β, Ccl2 mRNA was significantly decreased (p<0.0001) in ELAM+NMN-treated mice relative to NMN-treated mice (Fig 1i). While changes in expression levels were not detected in Cxcl10, the Ccl2 expression levels suggest that NMN treatment is associated with active signaling through the pro-inflammatory IL-1β pathway in the kidney, and that this is ameliorated by simultaneous ELAM treatment (Fig 1i).

In liver, Ccl2 mRNA was significantly upregulated only in untreated aged mice relative to young, with all aged treatment groups expressing varying levels that did not significantly differ from either untreated young or aged (Fig 1j). Like kidney, Cxcl10 mRNA expression did not yield any differences across age and treatment groups in the liver (Fig 1j). Taken together, these gene expression patterns suggest a kidney-specific inflammatory response to NMN supplementation that is modulated by the IL-1β pathway in aged animals and ameliorated by ELAM.

### IL-1𝛃 expression localizes to multiple kidney cell types in aged kidneys

We performed RNA in-situ hybridization (ISH) to localize which compartment(s) or cell(s) were responsible for elevated Il1β mRNA expression within the kidney. Consistent with other reports (33), expression of Il1β was sparse, but we saw instances of RNA-ISH positive signal detected in proximal and distal tubules, the interstitial space and the glomerulus (Fig 1k). This confirms that the tubule cells themselves are signaling for renal inflammation and that this is not solely driven by lymphocyte infiltrate.

To confirm proximal tubule damage, we measured mRNA for Kidney Injury Marker 1 (Kim-1), a marker of proximal tubule injury (34). Although the data had considerable variability, Kim-1 expression was only significantly upregulated in the kidneys of NMN-treated aged mice compared to young untreated mice (mean relative to young fold change: 21.06+/- SEM 6.05, p=0.019) by qRT-PCR. (Fig 1l). Kim-1 was also analyzed by RNA-ISH and demonstrated that expression was dispersed throughout the kidneys and was localized beyond obvious necrotic lesions. (Fig 1m).

The presence of IL-1β, a cytokine, in the proximal tubules and collecting duct cells in mice has been reported to induce mRNA expression of Lipocalin-2 (Lcn2/NGAL), a marker of distal tubule and collecting duct injury (28, 35). Therefore, we analyzed Lcn2 by RNA-ISH and expectedly found expression indicating that distal tubule injury was also present (Fig 1m). Overall, signal penetration with RNA-ISH was not consistent across individual sections and it was unclear whether this was due to variation in fixation or inherent RNA damage. Therefore, to avoid misinterpreting negative results, analysis of RNA-ISH is qualitative rather than quantitative.

### Age-specific changes to circulating metabolites in response to mitochondrial interventions suggests NMN driven uremia in aged mice

Both NMN and ELAM impact mitochondrial function, thus altering metabolic processes. Therefore, we analyzed the metabolic profiles of serum from this mice cohort and serum from an additional cohort of young mice (six-months-old at sacrifice). Young mice were treated identically to the aged cohort: eight weeks with either ELAM at 3mg/kg/day through an osmotic minipump, or NMN at 300mg/kg/day in the drinking water, or untreated control. Combined treatment was not studied in the young cohort. Using steady state metabolomics, we identified 46 metabolites in positive mode and 227 metabolites in negative mode. Although global metabolic analysis on the negative ion mode was conducted (Supplemental Fig 2), to best capture compounds closely related to the biosynthesis and structure of NAD^+^ we focused our analysis on those identified through the positive ion mode. To account for potential variation in loading between samples, we performed log transformation and mean centering of imputed values of the 19 metabolites determined to be significantly different between groups (Fig 3b-t).

To identify metabolites with effects of age, NMN and/or ELAM treatment as well as age-dependent effects of treatment, we ran an ANOVA (Methods, Fig 2a, Supplemental Fig 3). Nineteen distinct metabolites were identified as having significant effects: 15 associated with age, 11 associated with NMN, and one associated with ELAM, with some overlap (Fig 2a, Supplemental Table 1). By hierarchical clustering analysis of the significant metabolites associated with each effect (age, NMN or ELAM), the most striking pattern was the distinctive profile of NMN supplementation in both NMN- and ELAM+NMN- treated aged mice (Fig 2b). There was also a clear age-specific metabolic pattern that differed between young and aged mice independent of treatment (Fig 2b).

**Figure 2.**
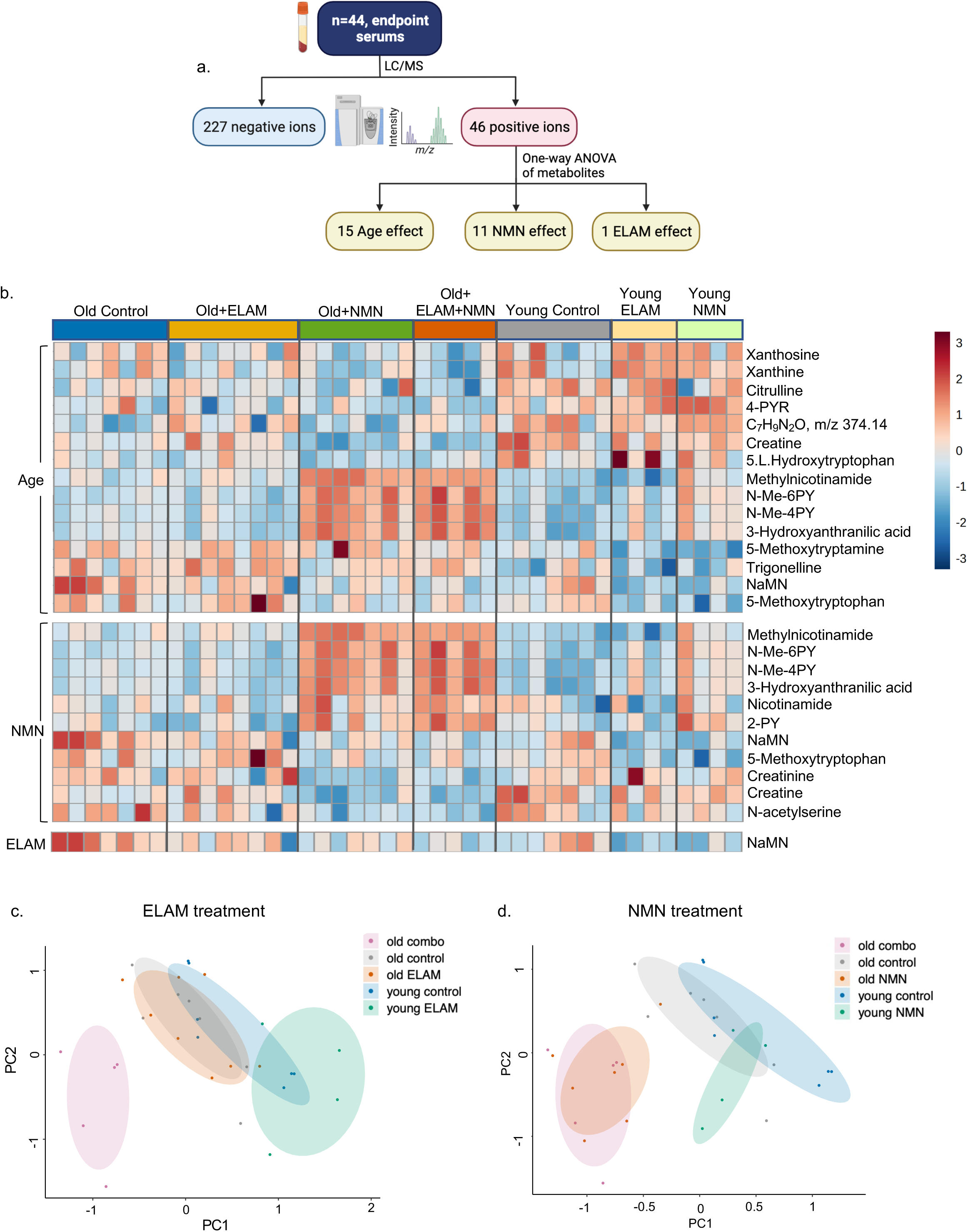
Metabolomic analyses reveals driving NMN signature. **a.** Flowchart of serum metabolome analysis. **b.** Hierarchical cluster analysis of metabolites with significant effects from ANOVA and a false discovery rate (FDR) of <5%. The top set of metabolites are significantly affected by age, the middle set are significantly affected by NMN, and the bottom metabolite is significantly affected by ELAM. Values shown are Z-scores in units of standard deviation. **c.** Principal Component Analysis (PCA) of young and aged, untreated (control) and NMN-treated animals, in addition to aged ELAM+NMN- (combo) treated animals. The ellipses are 60% confidence intervals for each group. **d.** PCA of young and aged control and ELAM-treated animals, in addition to aged combo treated animals. The ellipses are 60% confidence intervals for each group.

Principal component analysis (PCA) of the serum metabolome separated NMN-treated animals, in either age group, from age-matched controls (Fig 2c). Additionally, the clustering of aged ELAM+NMN treated animals overlapped with that of NMN treated animals (Fig 2c) but segregated completely from all ELAM-treated animals (Fig 2c & d), indicating that NMN is the primary driver of metabolic effects within the combined intervention. With respect to ELAM, the PCA plot showed that young ELAM treated mice segregated from untreated young mice, but that aged ELAM-treated did not segregate from untreated aged controls (Fig 2d). This finding was unexpected as previous studies from our group and others have not found a significant effect of ELAM on young adult animals (18, 19).

By one-way ANOVA the sera metabolites that differed most notably between groups, with significantly higher levels of detection in NMN- and NMN+ELAM-treated aged mice compared to aged control mice, were methylnicotinamide (MNA), N-methyl-6PY (N-Me-6PY), N-methyl-4PY (N-Me-4PY) and 3-hydroxyanthranilic acid (3-HAA) (Fig 3b, c, d, h). Although young NMN treated animals also had significantly increased levels of these metabolites compared to young untreated animals, the values were ultimately lower than those of aged NMN treated animals (Fig 3b, c, d, h), particularly for MNA which reaches a significantly higher value in the aged NMN treated mice compared to the young NMN-treated mice (Fig 3b). Levels of 2PY rose in both NMN-only and combined treatment groups, but only reached significance in the combined group relative to ELAM treated or untreated aged animals (Fig 3e). Fewer metabolic changes were detected in response to ELAM treatment alone (Fig 3f, i, j, n, s). It is important to note that N-Me-6PY is often referred to in the literature as 2-PY or N-Me-2PY. In untargeted LCMS, it has been recently reported that N-Me-2-PY and N-Me-4-PY tend to co-elute unless the elution method is specifically optimized to separate them. However, this optimization can compromise the separation of other catabolites. As a result, our N-Me-4-PY contains a mixture of both N-Me-2-PY and N-Me-4-PY (36).

**Figure 3.**
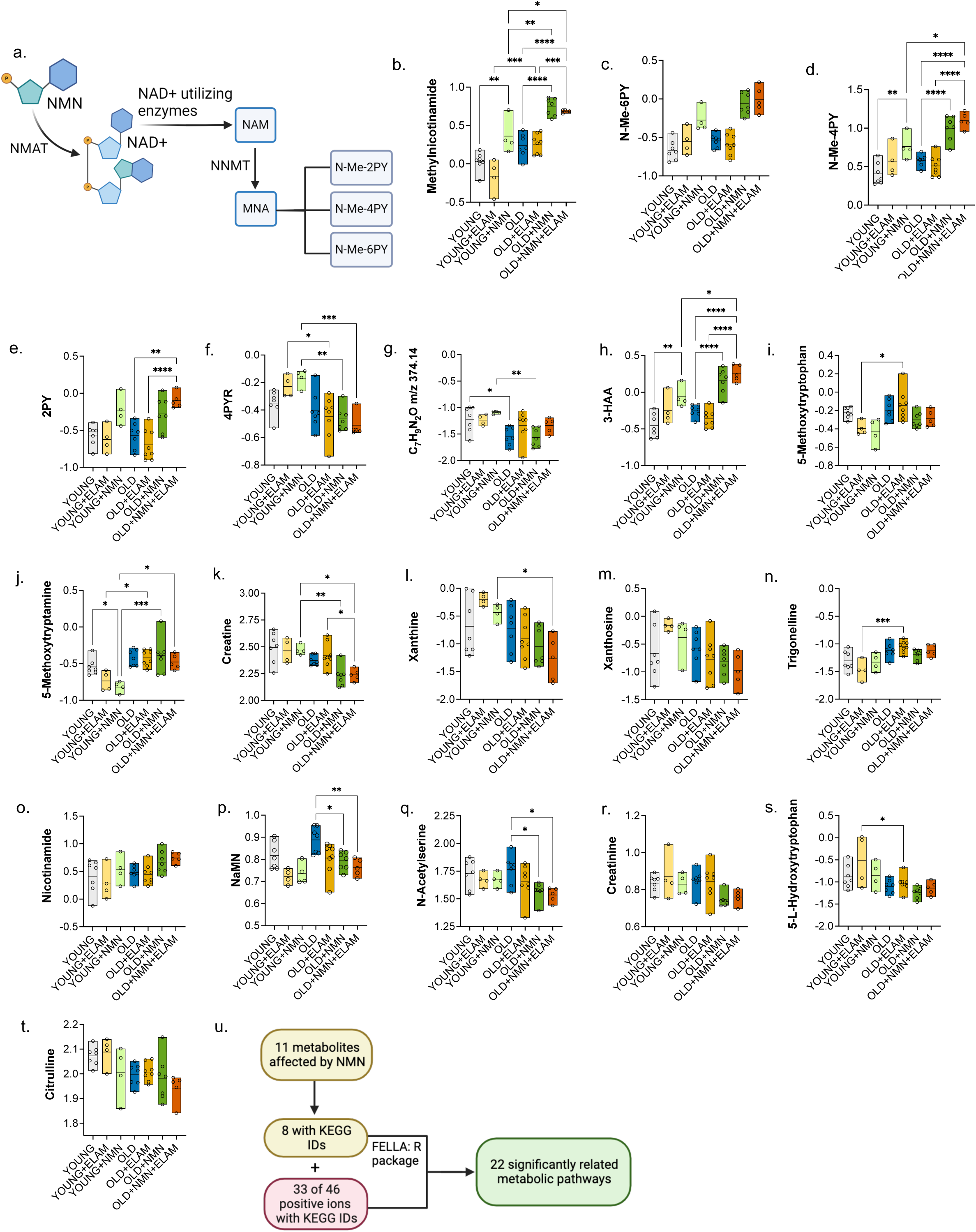
Significant age-associated variation in metabolite levels with ELAM and NMN supplementation. **a.** Schematic depicting pathway of metabolites and catabolites derived from NMN. **b-t.** The 19 metabolites identified to have significant variation due to age, NMN or ELAM by targeted one-way ANOVA (Fig 2b). To account for loading variation, values have been log- transformed, scaled by sample, and imputed for any missing data due to detection failure (n=8 values were imputed). Metabolites: **b.** Methylnicotinamide **c.** N-methyl-6-pyridone-5-carboxamide* (N-Me-6PY, 153.0658 m/z) **d.** N-methyl-4-pyridone-5-carboxamide (N-Me-4PY) **e.** N-methyl-2-pyridone-5-carboxamide* (2PY, 139.0501 m/z) **f.** 4-Pyridone-3-carboxamide-1-β-D-ribonucleoside (4PYR) **g.** C_7_H_9_N_2_O m/z 374.14 **h.** 3-Hydroxyanthranilic Acid (3-HAA) **i.** 5-Methoxytryptophan **j.** 5-Methoxytryptamine **k.** Creatine **l.** Xanthine **m.** Xanthosine **n**. Trigonelline (N-methyl-nicotinic acid,N-CH3-NA) **o.** Nicotinamide **p.** Nicotinate ribonucleotide (NaNM) **q.** N-Acetylserine **r.** Creatinine **s.** 5-L-Hydroxytryptophan **t.** Citrulline **u.** Flowchart depicting methodology for selected input compounds for pathway analysis. *Denotes compounds with same long-form nomenclature but distinguishable as separate compounds by M/S retention time and mass-charge (m/z) ratio.

Our findings were congruent with a previous human study of NMN supplementation that similarly reported increased circulating levels of MNA and 2-PY/4-PY in the serum of overweight, prediabetic, postmenopausal women who were administered NMN at 250 mg/day for 10 weeks (37). NMN is a precursor for the biosynthesis of NAD^+^ through the salvage pathway, which leads to metabolites via nicotinamide (NAM) methylation including MNA, N-Me-2-PY, N-Me-4-PY, and N-Me-6-PY. Similarly, 3-HAA is an intermediate of the kynurenine pathway of de-novo NAD^+^ synthesis, (Fig 3a) (36, 38, 39). Thus, these metabolites, related to catabolism of NAD^+^, are naturally occurring as a result of NAD^+^ synthesis and are found at detectable levels in young and healthy individuals; however, at elevated levels, they are considered uremic toxins (40, 41), which accumulate in the serum and urine of individuals with chronic kidney disease or otherwise impaired kidney function (39, 42–45). For example, one study identified the metabolome of urine from 82 injured combat soldiers, 33 who presented with acute kidney injury (AKI), and found MNA to be most associated with adverse outcomes of AKI and a predictive biomarker for mortality and renal replacement therapy (45).

To investigate the cellular and biological pathways that might underlie NMN-induced kidney inflammation and injury, we performed pathway analysis of the metabolites significantly associated with NMN (Methods, Fig 3u, Supplemental Table 2). Twenty-two biological pathways were enriched by the NMN-associated compounds, with an empirical P<0.01 (Table 1). When sorted by descending P value, the top ten pathways included broad signaling pathways with known mitochondrial axes such as apoptosis and necroptosis (46), and FoxO-signaling (47) (Fig 3u, Table 1). We also identified pathways with strong inflammatory association, such as complement and coagulation cascades (48) and the polycomb repressive complex (49).

**Table 1.** Ten pathways that are most significantly related to input metabolites that have significant variation due to NMN analyzed through the FELLA R-package using KEGG IDs. FDR-adjusted experimental p-scores listed, threshold p<0.01.

| KEGG ID | Pathway Name | p-score, <0.01 |
| --- | --- | --- |
| mmu03083 | Polycomb repressive complex | $1.84 \times 10^{-4}$ |
| mmu05322 | Systemic lupus erythematosus | $2.84 \times 10^{-4}$ |
| mmu04217 | Necroptosis | $7.84 \times 10^{-4}$ |
| mmu04211 | Longevity regulating pathway | $9.84 \times 10^{-4}$ |
| mmu05012 | Parkinson disease | $1.18 \times 10^{-3}$ |
| mmu05016 | Huntington disease | $3.28 \times 10^{-3}$ |
| mmu04210 | Apoptosis | $3.48 \times 10^{-3}$ |
| mmu05202 | Transcriptional misregulation in cancer | $3.68 \times 10^{-3}$ |
| mmu04068 | FoxO signaling pathway | $5.38 \times 10^{-3}$ |
| mmu04610 | Complement and coagulation cascades | $5.68 \times 10^{-3}$ |

Although the NMN- and ELAM+NMN- treated combined aged groups differed significantly in IL-1β and Ccl2 mRNA expression in the kidney (Fig 1a, i), the metabolic profiles were nearly identical (Fig 2c), suggesting that the mechanism of ELAM in a protective role against NMN-induced inflammatory signaling in the aged kidney does not involve decreasing levels of potential uremic metabolites. Collectively, these data show that accumulation of catabolites from NMN metabolism differs between aged versus young animals, with preexisting sera levels of some metabolites in aged animals increasing further in the context of NMN treatment.

### Peak and subsequent decrease in excretion of MNA and N-Me-4PY in urine shortly following NMN treatment suggests metabolite retention with long-term supplementation

Because sera were only collected at endpoint, we collected weekly spot urines from an additional cohort of aged untreated (n=4), NMN treated (n=6) and ELAM+NMN (n=5) treated animals for the duration of the 8-week treatment to better understand the longitudinal effects of ELAM and NMN supplementation on metabolic excretion. By two-way ANOVA there was a significant increase in the urine excretion of MNA (p<0.0001) and N-Me-4PY (p<0.0001) between ELAM+NMN treated and untreated aged mice two days after NMN supplementation was initiated (Fig 4a, b).

**Figure 4.**
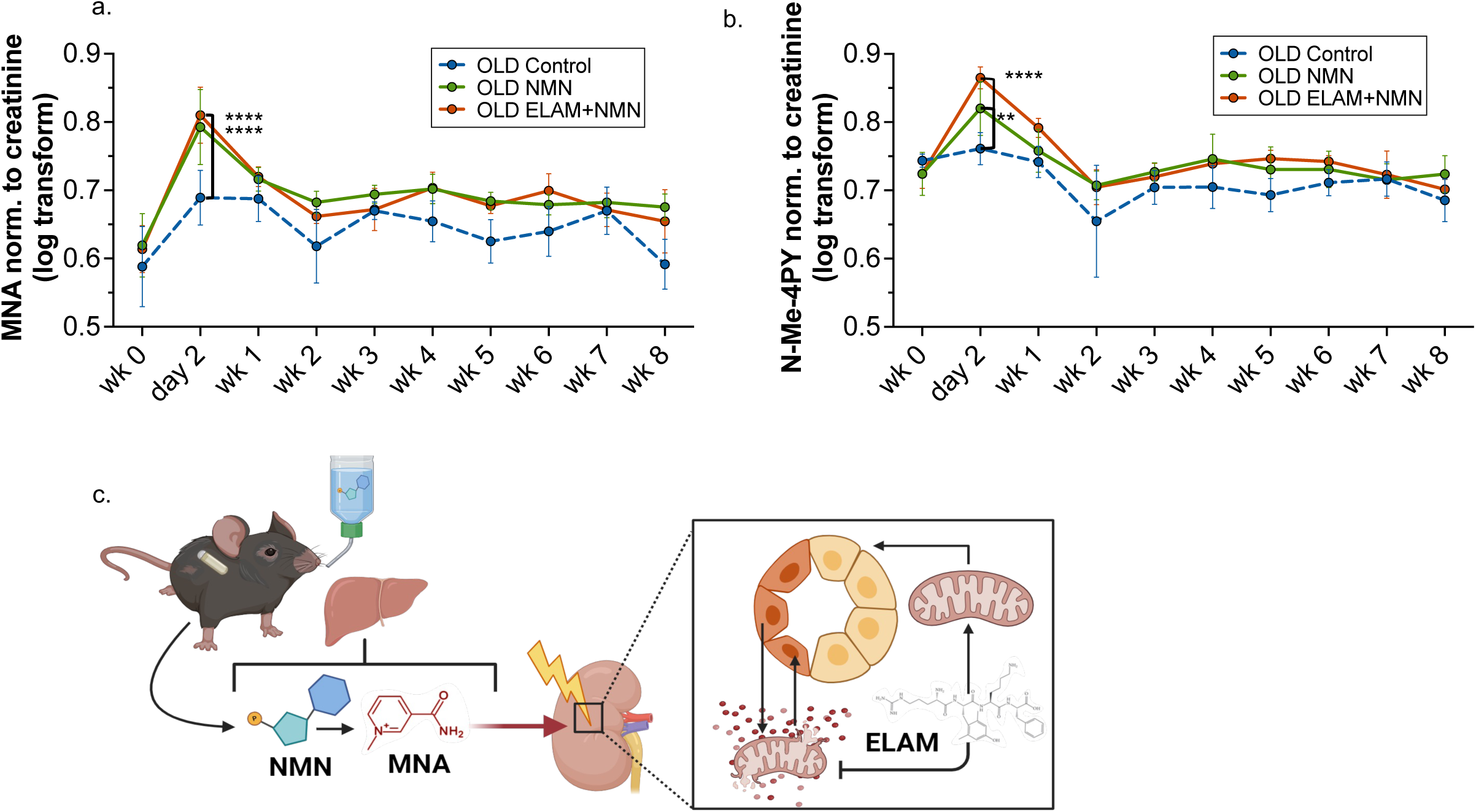
**a&b.** Log transform and mean centered values of metabolites (normalized to creatinine levels) in the weekly spot urines throughout 8 weeks of treatment. **** denotes p<0.0001, ** denotes p<0.01. **a.** Methylnicotinamide (MNA) **b.** N-methyl-4-pyridone-5-carboxamide (N-Me-4PY) **c**. Schematic of suspected mechanism of age-specific renal consequence of NMN supplementation and associated rescue from simultaneous ELAM supplementation.

Exclusively NMN-treated mice also had significantly higher excretion levels of MNA (p<0.0001) and N-Me-4PY (p=0.005) compared to untreated animals at day 2 (Fig 4a, b). This peak in urine excretion for NMN and combined treated animals is followed by a decrease in the detection of urinary of MNA and N-Me-4PY until week 2, when the values are closer to that of urines from untreated aged animals (Fig 4a, b). Coupled with the previous data indicating higher circulating levels of MNA and N-Me-4PY following 8 weeks of NMN supplementation in aged animals compared to untreated aged animals (Fig 3b, d), this suggests that efficient renal filtration of these NMN-derived catabolites wanes after two weeks of NMN supplementation possibly leading to a build-up of toxic catabolites in sera or other organ systems.

## Discussion

Many studies have reported that NMN is beneficial in boosting NAD^+^ levels in the context of age or disease, but few have focused on the kidney at late age (50–55). Our findings demonstrate that NMN supplementation aggravates preexisting inflammatory signaling in the kidneys of aged mice. Our results suggest that unlike heart and liver, the aged kidney may be uniquely susceptible to injury by excessive supplemental NMN, likely through the processing of uremic catabolites produced downstream of NMN metabolism. Previous studies focused on NMN supplementation in the kidney uniformly reported beneficial effects; however, all used acute toxic challenges or disease models rather than normative aging. Three separate studies demonstrated protective effects in the kidney tubule with short-term NMN supplementation against AKI in 2-3-month-old (56–58) or 20-month-old mice (56). In these studies, AKI was induced either by cisplatin or ischemia reperfusion with 500 mg/kg of NMN administered by intraperitoneal (IP) injection for 3-4 days at the onset of injury. NMN treated mice with AKI injury showed reduction of proteinuria, kidney tubule damage and mitochondrial dysfunction. Separately, kidneys of 24-month-old C57Bl/6 mice treated with NMN by 500 mg/kg IP injections every other day for 4 weeks had increased expression of tubule peroxisome associated proteins to levels more similar to young mice than untreated aged mice (59). In the kidney glomerulus, Yasuda et. al. reported decreased albuminuria and foot process effacement in addition to increased levels of nicotinamide phosphoribosyl transferase (NAMPT) and sirtuin 1 (SIRT1) in kidneys of 2-month-old diabetic db/db male mice treated with IP injections of 500 mg/kg of NMN daily for 2 weeks (52). Using the same dosing and duration, Hasawega et. al. reported restored podocyte numbers and ameliorated histological damage evidenced by Periodic acid-Schiff (PAS) and WT-1 stains in 2-month-old male mice with adriamycin-induced focal segmental glomerulosclerosis (60).

More studies exploring NMN supplementation in the context of aging have focused on the functional effects in liver (61–63), heart (21, 53, 64, 65), brain (66, 67) and skeletal muscle (54, 61, 68), and have primarily reported beneficial effects. Of note, we previously analyzed hearts in the same cohort of mice presented in this current study and similarly found functional benefits with NMN, ELAM, and combined NMN+ELAM treatment (21). In the livers of these same animals, we also showed a trending reduction in inflammatory markers, Ccl2 and IL-1β following NMN supplementation (Fig 1b, j) (21). These results in heart, liver, and kidney from a single animal cohort, underscore a critical aspect of our current findings, which is the organ specificity of potential benefits or detriments of late-age NMN supplementation. Nacarelli et al. also found tissue-specific inflammatory signaling with NMN, reporting that daily IP injections of 500 mg/kg of NMN increased pancreatic inflammation and fibrosis in 2-month-old mice with pancreatic cancer as evidenced by increased mRNA expression of IL-1β, IL-6, p16, and immunohistochemistry for F4/80 positive macrophage tissue infiltration (69).

A final significant finding of our study was the age-specific accumulation of NMN catabolites. Our work did not distinguish between tissue sites of NMN processing, such as the liver versus kidney, and thus we cannot speculate on whether dysfunction in other aged tissues contributed to the accumulation of the purported uremic toxins. Because our study analyzed endpoint tissues at 8 weeks of treatment we also do not know when the IL-1β expression increased or if the enhanced IL-1β signaling would ultimately contribute to acceleration of nephrotoxic inflammation; however, several studies have shown that IL-1β signaling is indicative of worsening chronic kidney disease and acute kidney injury in both mice and humans (27, 33, 70, 71). One indication to the age-specific differences was demonstrated by the combined ELAM+NMN treatment group, in which we detected a significant reduction in NMN induction of IL-1β inflammatory signaling despite no significant rescue of histopathology or metabolite expression between ELAM+NMN versus NMN treatment. This suggests that the damage from NMN treatment is distinct from mitochondrial rescue, but that underlying mitochondrial dysfunction from aging may contribute to exacerbation of inflammatory signaling. The mechanism through which ELAM may be reducing IL-1β signaling in the presence of uremia remains unclear; however, previous studies have shown that ELAM treatment reduces IL-1β signaling in an acute kidney injury model in mice (72) as well as in a diet-induced model of metabolic syndrome in pigs (73).

In our study, treatment with ELAM resulted in a distinct metabolic profile in endpoint sera from young mice. Although not reaching significance by one-way ANOVA, we observed a trend where ELAM-only treatment in aged mice restored the levels of some metabolites closer to the levels measured in young mice. This contrasted with the NMN-specific profiles of aged mice, which increased detection of some metabolites to levels significantly higher than were observed in young groups. Moreover, the pathways engaged by the two interventions were distinct, as ELAM increased circulating levels of 5-methoxytryptophan and creatine (Fig 3i, k), whereas NMN increased MNA, N-Me-6PY, N-Me-4PY and 3-HAA (Fig 3b, c, d, h). The definitive roles of these metabolites in inflammation, aging-related mitochondrial dysfunction, and energetics are still unclear and the literature is split on whether increasing circulating levels are indicative of benefit or harm for most metabolites. A recent study by Itoh et al used unilateral ureter obstruction with supplementation of NMN catabolite, N-Me-2PY, at 300 mg/kg/day in mice and demonstrated anti-fibrotic effects (74). Unfortunately, the age of the mouse cohort was not reported, making conclusions about age susceptibility unavailable. In heart, levels of 2PY and 4PY, which are also catabolites of NMN metabolism, have recently been shown to positively correlate to increased risk of major adverse cardiac events in patients from increased vascular inflammation (75). The same study showed that in young mice, exogenous treatment with 4PY, but not 2PY, increased vascular inflammation through increased vascular adhesion molecule (VCAM-1) and leukocytes to endothelial adhesion. Based on our results, we suspect that the physiologic balance of these metabolites is the critical measurement and that a significant increase over young adult levels, as we saw in NMN-treated aged mice, appears to indicate detriment rather than benefit.

While this study focuses on an understudied aspect of NMN supplementation tissue (kidney) and aging, (the 26-month mice are equivalent to a 79-year-old in humans), our study shows unexpected results that highlight remaining gaps of knowledge. First, although this study focused on male mice, we expect there to be sex-specific differences in NMN processing such as that reported by van der Velpen et. al. with upregulation of tryptophan-related metabolites in female NMN-treated mice compared with male NMN-treated mice with Alzheimer’s disease (76).

Second, while we saw a significant increase in inflammatory chemokine levels in NMN-treated aged mice, F4/80 histology of macrophage infiltration did not change with treatment (Supplemental Fig 1). However, this represents just a single inflammatory cell population and may not have fully captured infiltration of leukocytes. Third, this study lacked longitudinal sera and measures of continuous kidney injury, which limited our understanding of when the NMN-induced damage occurred during the 8-week treatment. Fourth, despite identifying possible pathways connected to the metabolic signature of NMN, this study did not further assess the implications of mitochondrial dysfunction interventions on entire metabolic networks (Table 1). Fifth, it is important to note that mice and humans possess differing levels and forms of some NAD/Me- NAM metabolic enzymes which can affect the impact of NMN or other NAD^+^ compounds in a species-specific manner (77). Lastly, if the dosing of NMN in the drinking water of our study were allometrically scaled for human consumption, it would be approximately 1.5 gram/day (78). This is higher than what has been used in human clinical trials (100-900 mg/day) (61, 79–81), but is consistent with at least 6 other studies using oral NMN delivery in mice in a range between 300-500 mg/kg/body weight (82). On the other hand, the ELAM dosing, if scaled, was less than half the dose used for human subjects across clinical trials (83). Furthermore, this dosing of NMN is still a more realistic model for the potential for excessive self-dosing, which is enabled by the current availability of many NAD^+^ precursors as over-the-counter supplements.

Although approval for NMN to be sold as an openly available dietary supplement was revoked by the United States Food and Drug Administration in October of 2022, this change was made in response to an investigational new drug application intended to place a formulation of NMN into clinical trials as a ‘new drug’ (84). We encourage future studies, particularly clinical trials, to expand the reach of our current work, by ensuring that kidney function is included as an important safety measure. We suggest investigating sex specificity, a broader set of inflammatory markers and to continually assess uremic metabolite levels in urine and serum compared to baseline kidney function with analysis stratified by age. In addition, to best replicate the potential of chronic use in humans, future experiments should expand the duration of treatment. Overall, this study illustrates the difficulty in assessing the systemic efficacy of aging interventions with various regimens, ages, and tissue effects. However, considering the availability of NAD^+^ precursors for the public, the inconsistency in risk and benefit across tissues continues to be concerning (51, 53, 64, 67, 69, 85). The limited regulations around consumption of supplements intended to boost NAD^+^ levels underscore the urgent need for increased caution and further studies in the use of such supplements for public safety.

## Materials and Methods

### Animals and Treatment

Male C57BL/6J mice, from the National Institute on Aging Charles River colony were housed at 20°C in the AAALAC accredited University of Washington Medical Center Facilities upon approval from the Office of Animal Welfare. 24 months of age mice were divided randomly into four groups. 1) ELAM was provided through a material transfer agreement between the University of Washington and Stealth BioTherapeutics (Needham, MA). ELAM was delivered through automatic osmotic minipumps (Alzet, Cupertino, CA) at 3 mg/kg body weight per day, with new minipumps substituted after 4 weeks to continue treatment. 2) As reported, individual cage water consumption was measured for one week prior to start of experimentation (21). NMN was fed through the drinking water with a concentration based on each cage’s measured water consumption rate and mean mouse body weight amounting to 300 mg/kg body weight per day as the only source of hydration and was replaced after 3-7 days. 3) ELAM+NMN were simultaneously given through combined minipump and drinking water. 4) Untreated mice were housed with ELAM osmotic mini-pump mice. After 8 weeks of treatment, mice were euthanized, and tissues were collected. Untreated mice were sacrificed at 6- and 26-months-of-age to serve as age-matched control groups. Results from this same mouse cohort were previously published (21).

### Quantitative Real Time PCR

Total RNA was extracted using the RNeasy kit (Qiagen-Hilden, Germany: Ref#74104). Reverse transcription was completed using QuantiNovaTM RT kit (Qiagen-Hilden, Germany: Ref#205413) according to manufacturer’s guidelines. Quantitative Real-Time PCR was conducted using Taq Man Gene Expression Master Mix (Applied Biosystems-Thermo Scientific-Vilnius, Lithuania: catalog #4444557) in the Rotor-Gene Q real-time PCR cycler machine (Qiagen, catalog #) at a 10 µL reaction volume. The following pre-made *Mus musculus* primers (Applied Biosystems, catalog #4331182) were used in the qRT-PCR panel: Actb, Cdkn2a (p16^INK4A^), Cdkn1a (p21), Trp53 (p53), Lmnb1, IL-1β, Hmgb1, Havcr1 (Kim-1), Ccl2, Cxcl10 and Bcl3. Expanded information is available in supplemental methods.

### RNA In-Situ Hybridization

RNA In-Situ Hybridization (RNA-ISH) RED Detection (ACD Bio RNAscope 2.5 HD Reagent Kit-RED assay; catalog #322350) was performed on formalin-fixed paraffin-embedded kidney tissue. The same paraffin blocks were used for both Immunohistochemistry and RNA-ISH. Slides (n>4 for each treatment group) were hybridized with either the IL-1β probe (ACD Bio RNA scope Probe, Mm-Il1b; catalog #316891), Havcr1 probe (ACD Bio RNA scope Probe, Mm-Havcr1; catalog #472551), Lcn2 probe (ACD Bio RNA scope Probe -Mm-Lcn2; catalog #313971) or the positive housekeeping control peptidylprolyl isomerase B (P Ppib, ACD Bio RNA scope, Mm-Ppib; catalog #313911) with protease retrieval and hybridization per manufacturer protocols.

### NAD^+^ Metabolomics

Serum was thawed on ice before adding 13.5 μL of −80°C 100% methanol per 1 μL of serum. Samples were vortexed and incubated on dry ice for 10 minutes, followed by centrifugation at 16,000 g for 25 minutes. The supernatant diluted to 80:20 methanol: water. Urine was thawed on ice and 10 μL was extracted with 90 μL of −80°C 100% methanol, vortexed then centrifuged at 16,000 g for 25 minutes. Supernatants were collected and the pellet further extracted using 100 μL of 40:40:20 acetonitrile:methanol:water. Mixture was vortexted and then centrifuged at 16,000 g for 25 minutes. The resulting supernatant was combined with the original supernatant, mixed, and centrifuged at 16,000 g for 25 minutes before by liquid chromatography coupled to a mass spectrometer (LC-MS) analysis.

Extracts were analyzed within 24 hours by LC-MS. The LC–MS method was based on hydrophilic interaction chromatography (HILIC) coupled to the Orbitrap Exploris 240 mass spectrometer (Thermo Scientific) (86). The LC separation was performed on a XBridge BEH Amide column (2.1 x 150 mm, 3.5 μm particle size, Waters, Milford, MA). For the detection of metabolites, the mass spectrometer was operated in both negative and positive ion mode. Data were analyzed via the EL-MAVEN software version 12. Expanded information in supplemental methods.

### Statistical Analysis

GraphPad Prism 9.0 (La Jolla, CA) was used for statistical analysis of qRT-PCR data. Two-way ANOVA was used for Ccl2, Cxcl10, Cdkn1a, Cdkn2a and Trp53 in kidney and liver. One-way ANOVA was used for Lnmb1, IL-1β in kidney and liver, Bcl3 in liver and Kim-1 in kidney. All qRT-PCR results are plotted as means +/- SEM with p-values listed.

For metabolomics, R-studio (Boston, MA) and R version 4.3.2 were used to log-transform LC-MS peak intensities, scale by sample, and impute missing metabolite values for metabolites with ≤4 missing values (n=9 total values imputed by 10-nearest neighbor imputation) (87). Principal components analysis (PCA) was performed and one ELAM+NMN treated sample was detected as an outlier, having a Mahalanobis distance over the first two PCs of greater than three standard deviations and removed from further analysis. For univariate analysis, we ran ANOVA on each metabolite in a linear model where main effects of mouse age, NMN treatment, ELAM treatment, as well as interaction terms for NMN x age, and ELAM x age were simultaneously fit to the data. In this model, NMN and ELAM treatment effects were both fit to the data from mice with the NMN+ELAM combination treatment. We controlled the false discovery rate (88). The ‘pheatmap’ R-package was used for hierarchical clustering analysis of Z-scores in units of standard deviation. PCA plots of PC1 and PC2 were created with a 60% confidence interval for each group. We used GraphPad Prism to get statistics for individual metabolites from a one-way ANOVA adjusted for multiple comparisons using the Šidák correction. Significance of pairwise comparisons denoted with * p≤0.05, ** p≤0.01, ***p ≤0.001 and **** p<0.0001.

Pathway enrichment analysis was done in the FELLA R-package on the metabolites with NMN effects using the remaining measured metabolites as background (89). Only metabolites with a corresponding ID in the KEGG database can be analyzed (90). Of the 11 metabolites with NMN effects, eight had KEGG IDs, and likewise for 33 of the remaining 35 background compounds (Fig 3u) (89, 90). Enrichment of pathways was measured using the diffusion method followed by 10,000 randomizations of the input compounds among the background compounds to derive empirical FDR-adjusted P values for each pathway (p-score). MetaboAnalyst was used for hierarchical clustering of heatmap that included all metabolites (Supplemental Fig 2) using 1-Pearson correlation as a distance measure and Ward clustering method.

Raw LC-MS values for metabolomics of spot urines were log-transformed, mean-centered using R-script and normalized to creatinine values of the same sample (Fig 4 a, b). GraphPad Prism was used for statistical analysis. Two-way ANOVA was used with significance of multiple comparisons denoted with * p≤0.05, ** p≤0.01, ***p ≤0.001 and **** p<0.0001 in the figure.

## Acknowledgments

We thank Dr. Shin Ichiro-Imai for supplying NMN received from Mirai Lab Bioscience, Tokyo, Japan. We would also like to acknowledge the Huck Institutes’ Metabolomics Core Facility (RRID:SCR_023864) for use of the OE 240 LCMS and Sergei Koshkin for helpful discussions on sample preparation and analysis. We thank Dr. Benjamin Harrison for assistance in metabolomic data analysis and presentation. We thank Dr. Marie Migaud, professor of Pharmacology at University of South Alabama for her extremely helpful guidance on accurate metabolite classification. Some figures used Biorender.com for the platform to create the schematics in this manuscript. This work was funded by the National Institutes of Health: National Institute on Aging (K01AG062757 to MTS, P01AG001751 to PSR), the Howard Hughes Medical Institute Hanna H. Gray Fellows Program Faculty Phase (Grant# GT15655 awarded to MRM); and the Burroughs Welcome Fund PDEP Transition to Faculty (Grant# 1022604 awarded to MRM).

## Author Contributions

MTS guided and supervised the study. TAS, PK, CNM and MTS collected tissues and conducted experimentation. JW and PSR provided murine samples. JW, PK, CNM and MTS collected urines and were responsible for animal treatment and monitoring. PP, BCJ, TS and MRM conducted metabolomics experimentation. TAS, PP, BCJ, MRM, PK, BRH and MTS analyzed metabolomics. TAS and MTS wrote the initial draft of the manuscript with all authors contributing to the final version.

## Figures and Tables

**Supplemental Figure 1.**
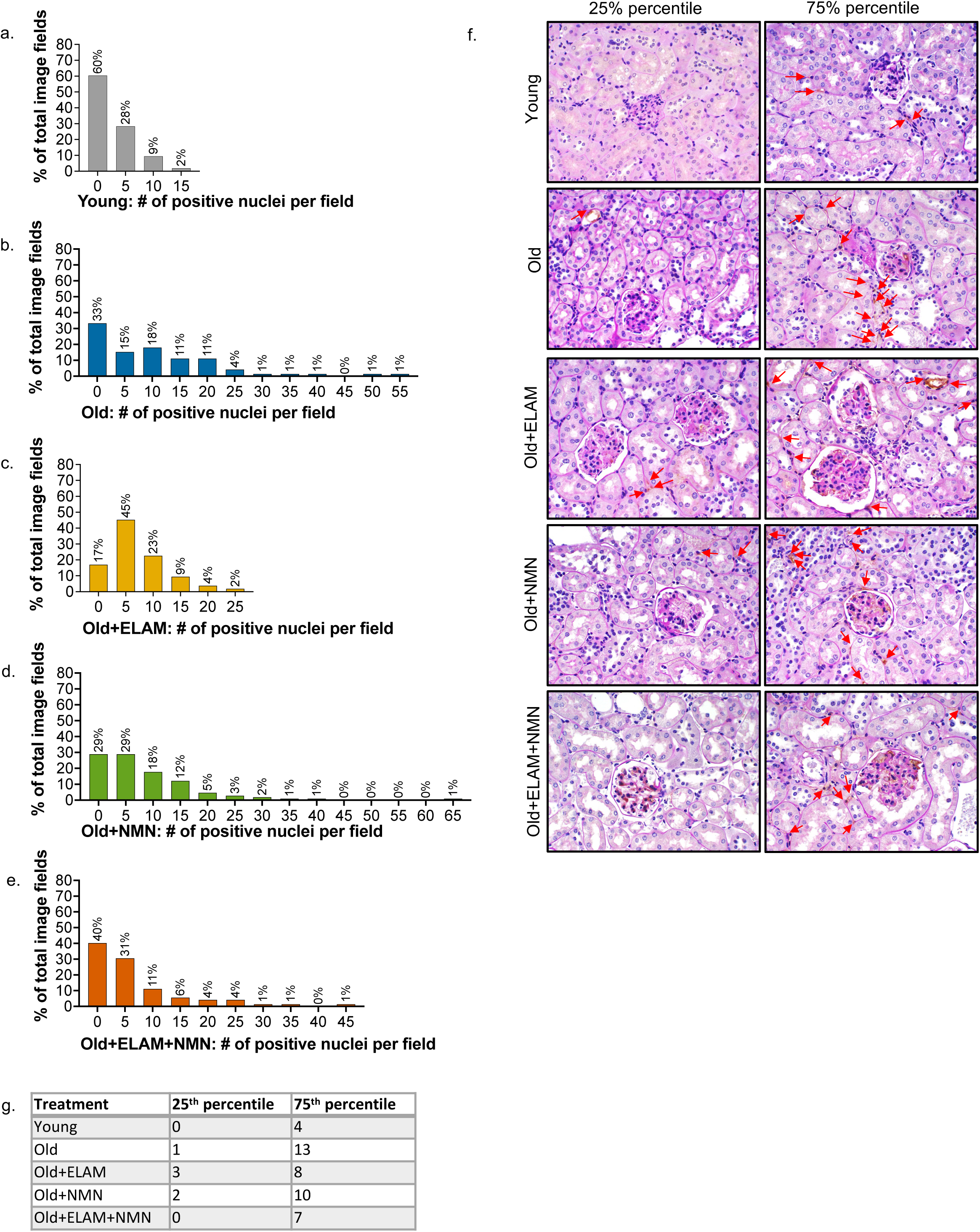
**a-e.** Frequency distribution graphs of number of F4/80+ positive nuclei per field. To assess monocytes and macrophage infiltration in the kidney, rat anti-mouse antibody for F4/80, Slides were counterstained using standard protocol of PAS. Eighteen image fields were taken per mouse. Values are binned with a bin size of 5. **f.** Representative images of the 25^th^ and 75^th^ percentile of number of positive nuclei in each treatment group (**g.)** with co-staining using PAS (Periodic acid-Schiff) and F4/80 IHC, a mouse macrophage marker. Positive nuclei shown with red arrows. **g.** Number of F4/80+ positive nuclei per field in the 25^th^ and 75^th^ percentile for each treatment group.

**Supplemental Figure 2.**
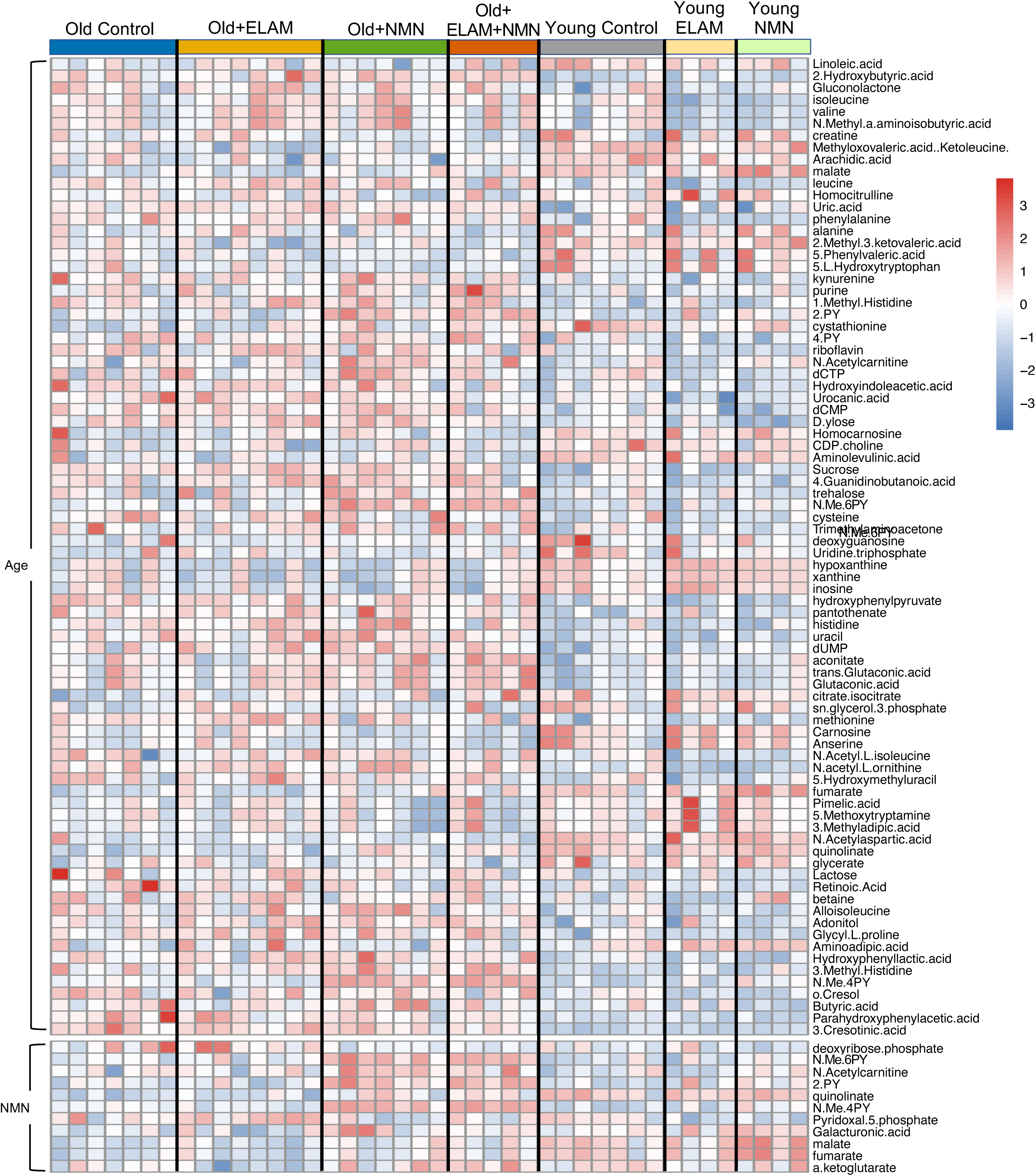
Hierarchical cluster analysis of metabolites with significant effects from ANOVA and a False Discovery Rate (FDR) of <5%. The variance in the top 82 metabolites were significantly affected by age, while the variance of the bottom 11 metabolites was significantly affected by NMN. The metabolome was extracted, and global analysis of the negative ion mode was obtained via LC-MS. Heatmap made in R-studio, values shown are Z-scores in units of standard deviation.

**Supplemental Figure 3.**
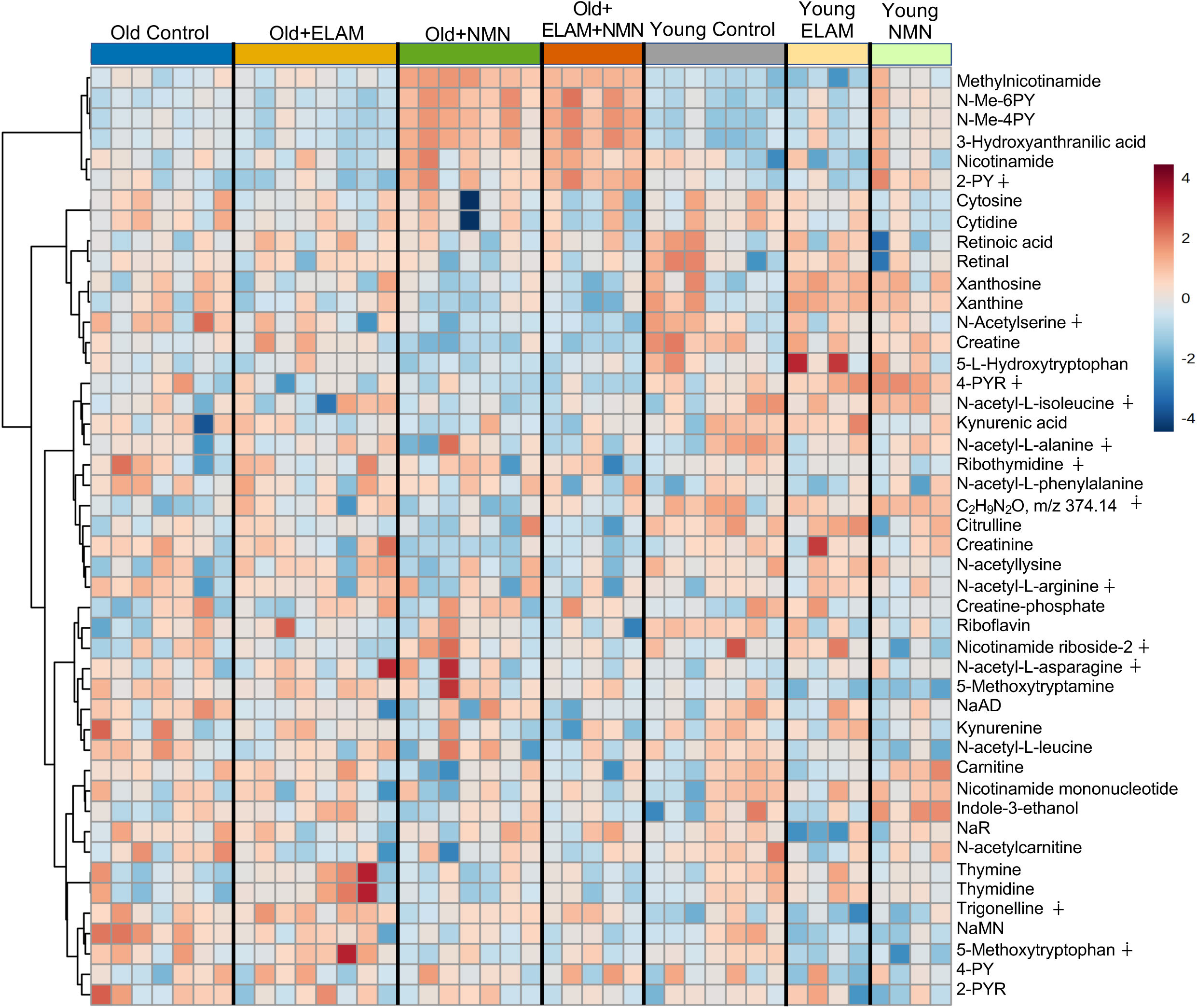
Hierarchical cluster analysis of all 46 metabolites in positive ion mode. Sample values were log-transformed and mean centered. The metabolome was extracted, and targeted analysis was obtained via LC-MS. Heatmap made in MetaboAnalyst. Values shown are Z-scores in units of standard deviation. (∔) indicates no associated KEGG ID.

**Supplemental Table 1.** The table includes the 19 metabolites that were distinct among those significantly affected by either age (15 metabolites), NMN (11 metabolites) or ELAM (1 metabolite) by one-way targeted ANOVA. n.f.= KEGG ID not found. (ϑ) indicates indistinguishable KEGG IDs. p-values were adjusted by False Discovery Rate <5%. Significant p-values <0.05 are denoted with an asterisk (*) and bolding.

| Metabolites | KEGG ID | Formula | m/z | Retention Time | Age (p-value) | All NMN (p-value) | All ELAM (p-value) |
| --- | --- | --- | --- | --- | --- | --- | --- |
| Methylnicotinamide | C02918 | C7H8N2O | 137.071 | 7.74 | **\*3.59 x 10<sup>-8</sup>** | **\*3.53 x 10<sup>-10</sup>** | 7.17 x 10<sup>-1</sup> |
| N-Me-6PY (θ) | C05842 | C7H8N2O2 | 153.066 | 3.64 | **\*5.18 x 10<sup>-4</sup>** | **\*1.19 x 10<sup>-10</sup>** | 8.80 x 10<sup>-1</sup> |
| N-Me-4PY | C05843 | C7H8N2O2 | 153.066 | 3.89 | **\*2.36 x 10<sup>-4</sup>** | **\*3.53 x 10<sup>-10</sup>** | 7.17 x 10<sup>-1</sup> |
| 2-PY (θ) | C05482 | C6H6N2O2 | 139.050 | 6.20 | 5.87 x 10<sup>-1</sup> | **\*7.02 x 10<sup>-7</sup>** | 9.56 x 10<sup>-1</sup> |
| 4-PYR | n.f. | C11H14N2O6 | 271.090 | 7.56 | **\*1.29 x 10<sup>-4</sup>** | 9.85 x 10<sup>-1</sup> | 9.56 x 10<sup>-1</sup> |
| C<sub>7</sub>H<sub>9</sub>N<sub>2</sub>O, m/z 374.14 | n.f. | C7H9N2O | 374.140 | 7.88 | **\*1.19 x 10<sup>-3</sup>** | 8.76 x 10<sup>-1</sup> | 3.48 x 10<sup>-1</sup> |
| 3-Hydroxyanthranilic acid | C00632 | C7H7NO3 | 154.050 | 3.88 | **\*4.59 x 10<sup>-4</sup>** | **\*3.53 x 10<sup>-10</sup>** | 7.17 x 10<sup>-1</sup> |
| 5-Methoxytryptophan | n.f. | C12H14N2O3 | 235.103 | 12.34 | **\*3.55 x 10<sup>-2</sup>** | **\*6.77 x 10<sup>-3</sup>** | 9.56 x 10<sup>-1</sup> |
| 5-Methoxytryptamine | C05659 | C11H14N2O | 191.122 | 7.47 | **\*9.31 x 10<sup>-5</sup>** | 3.18 x 10<sup>-1</sup> | 7.12 x 10<sup>-1</sup> |
| Creatine | C00300 | C4H9N3O2 | 132.077 | 7.28 | **\*2.36 x 10<sup>-4</sup>** | **\*6.66 x 10<sup>-3</sup>** | 9.56 x 10<sup>-1</sup> |
| Xanthine | C00385 | C5H4N4O2 | 153.041 | 6.36 | **\*2.16 x 10<sup>-3</sup>** | 2.56 x 10<sup>-1</sup> | 9.56 x 10<sup>-1</sup> |
| Xanthosine | C01762 | C10H12N4O6 | 285.083 | 6.36 | **\*3.54 x 10<sup>-2</sup>** | 4.85 x 10<sup>-1</sup> | 9.56 x 10<sup>-1</sup> |
| Trigonelline, N-CH3-NA | n.f. | C7H8NO2 | 139.060 | 6.65 | **\*9.31 x 10<sup>-5</sup>** | 4.85 x 10<sup>-1</sup> | 9.56 x 10<sup>-1</sup> |
| Nicotinamide | C00153 | C6H6N2O | 123.055 | 3.71 | 8.51 x 10<sup>-2</sup> | **\*1.85 x 10<sup>-2</sup>** | 9.56 x 10<sup>-1</sup> |
| NaMN | C01185 | C11H14NO9P | 336.053 | 5.06 | **\*4.86 x 10<sup>-2</sup>** | **\*1.42 x 10<sup>-2</sup>** | **\*2.55 x 10<sup>-2</sup>** |
| N-Acetylserine | n.f. | C5H9NO4 | 148.061 | 7.52 | 2.25 x 10<sup>-1</sup> | **\*1.85 x 10<sup>-2</sup>** | 4.67 x 10<sup>-1</sup> |
| Creatinine | C00791 | C4H7N3O | 114.066 | 4.97 | 1.75 x 10<sup>-1</sup> | **\*1.64 x 10<sup>-2</sup>** | 4.62 x 10<sup>-1</sup> |
| 5-L-Hydroxytryptophan | C01017 | C11H12N2O3 | 221.092 | 5.85 | **\*1.07 x 10<sup>-3</sup>** | 2.93 x 10<sup>-1</sup> | 9.11 x 10<sup>-1</sup> |
| Citrulline | C00327 | C6H13N3O3 | 176.103 | 7.60 | **\*4.18 x 10<sup>-3</sup>** | 9.28 x 10<sup>-2</sup> | 9.56 x 10<sup>-1</sup> |

## Supplementary Materials and Methods

### Quantitative Real Time PCR

In testing for housekeeping genes, the data showed variability in the upkeep of the Rn18s and Gapdh primers for the aged animals, with more consistent results seen with the β-Actin. PCR was initially denatured at 95°C for 10 minutes, followed by a cycle consisting of 95°C for 15 s and 60°C for 60 s for 40 cycles. The data was normalized to β-Actin levels and analyzed with the ΔΔ Ct method. RNA Integrity Number (RIN#) was confirmed by Tapestation, and any samples below the threshold were removed from the dataset and analysis. Additional outliers were removed according to the ROUT method with a threshold >2%.

### Immunohistochemistry

From the kidney, the anatomical left side was dissected, decapsulated and butterflied. From the liver, a section was taken from the anterior lobe. All tissue samples were kept in cassettes and fixed in buffered 4% paraformaldehyde overnight (∼12 hours) before transfer to 70% ethanol for storage in a 4°C refrigerator. The cassettes were then embedded in paraffin. Sections, 4 µm thick, were deparaffinized in xylene and rehydrated using a gradient of ethanol (70%-100% concentration solutions). To assess monocytes and macrophage infiltration in the kidney, rat anti-mouse antibody for F4/80, clone BM8 (Invitrogen, Carlsbad, CA) was used. Secondary reagents used for immunohistochemistry were ImmPRESS anti-rabbit horseradish peroxidase (HRP; Vector Laboratories, Burlingame, CA), ImmPRESS anti-goatHRP (Vector Laboratories), ImmPRESS anti-rat HRP, ImmPRESSanti-rat AP, MM HRP-Polymer kit (Biocare, Concord, CA), andPolink-1 HRP rat NM kit (Golden Bridge, Mukilteo, WA) followed by development with 3,3-diaminobenzidine. Slides were counterstained using standard protocol of PAS.

### Urinalysis

Spot urines were collected weekly from all mice throughout the 8-week treatment, beginning shortly before treatment administration. Proteinuria was quantified by albumin to creatinine ratio (ACR). Albumin and creatine concentrations were measured using the Albuwell M Elisa Kit (Exocell, Ethos Bioscience Ref#1011) and Creatinine Colorimetric Assay Kit (Cayman Chemical, Ref#500701), respectively. Six μL urine was diluted with 102 μL diluent for the albumin assay and 2.4 μL urine was diluted with 33.6 μL diluent for the creatinine assay, and both were analyzed according to manufacturer instructions. Albumin analysis was rerun if the coefficient of variation (CV) of replicate wells on the initial run exceeded 10%, with two runs being the maximum number completed for all three treatment and control groups.

### NAD^+^ Metabolomics

#### Serum Extraction

Serum was thawed on ice before adding 13.5 μL of −80°C 100% methanol per 1 μL of serum. Samples were vortexed and incubated on dry ice for 10 minutes, followed by centrifugation at 16,000 g for 25 minutes. The supernatant was then diluted to 80:20 methanol:water and used for LC-MS analysis.

#### Urine Extraction

Urine was thawed on ice, and 10 μL of urine was transferred to an Eppendorf tube. Then, 90 μL of −80°C 100% methanol was added for extraction. The urine samples were vortexed and centrifuged at 16,000 g for 25 minutes. The supernatants were carefully transferred to new Eppendorf tubes. The remaining pellet was further extracted using 100 μL of 40:40:20 acetonitrile:methanol:water. After vortexing, the mixture was centrifuged at 16,000 g for 25 minutes. The resulting supernatant was combined with the original supernatant collected, vigorously vortexed to ensure thorough mixing, and centrifuged at 16,000 g for 25 minutes before liquid chromatography-mass spectrometry (LC-MS) analysis.

#### LC-MS

Extracts were analyzed within 24 hours by LC-MS. The LC–MS method was based on hydrophilic interaction chromatography (HILIC) coupled to the Orbitrap Exploris 240 mass spectrometer (Thermo Scientific) (53). The LC separation was performed on an XBridge BEH Amide column (2.1 x 150 mm, 3.5 μm particle size, Waters, Milford, MA). Solvent A was 95%: 5% H2O: acetonitrile with 20 mM ammonium acetate and 20mM ammonium hydroxide, and solvent B was 90%: 10% acetonitrile: H2O with 20 mM ammonium acetate and 20mM ammonium hydroxide. The gradient was 0 min, 90% B; 2 min, 90% B; 3 min, 75% B; 5 min, 75% B; 6 min, 75% B; 7 min, 75% B; 8 min, 70% B; 9 min, 70% B; 10 min, 50% B; 12 min, 50% B; 13 min, 25% B; 14min, 25% B; 16 min, 0% B; 18 min, 0% B; 20 min, 0% B; 21 min, 90% B; 25 min, 90% B. The following parameters were maintained during the LC analysis: flow rate 150 mL/min, column temperature 25 °C, injection volume 5 µL and autosampler temperature 5 °C. For the detection of metabolites, the mass spectrometer was operated in both negative and positive ion mode. The following parameters were maintained during the MS analysis: resolution of 180,000 at m/z 200, automatic gain control (AGC) target at 3e6, maximum injection time of 30 ms and scan range of m/z 70-1000. Raw LC/MS data were converted to mzXML format using the command line “msconvert” utility (54). Data were analyzed via the EL-MAVEN software version 12.

